# Spinal Cord Ultrasound Stimulation Modulates Corticospinal Excitability

**DOI:** 10.1101/2025.01.13.632757

**Authors:** Lin Hou, Yuming Lei

**Affiliations:** Program of Motor Neuroscience Department of Kinesiology & Sport Management Texas A&M University, College Station, TX, 77843

**Author notes:** Correspondence to: Yuming Lei, PhD. Department of Kinesiology & Sport Management Texas A&M University.

**Keywords:** Ultrasound stimulation, Spinal cord, Corticospinal excitability, Transcranial magnetic stimulation, Human

## Abstract

**Background:** Low-intensity focused ultrasound (LIFU) offers superior tissue penetration and enables precise neuromodulation of cortical and subcortical circuits. However, its effects on neural activity in the human spinal cord remain largely unexplored.

**Objective:** To investigate the effects of LIFU on spinal cord neuromodulation under varying conditions of intensity (spatial-peak pulse-average intensity, I_SPPA_), duty cycle (DC), and pulse repetition frequency (PRF).

**Methods:** Thirty-six healthy human volunteers participated in the study. A 500 kHz ultrasound transducer with a focal depth exceeding 100 mm was used to target the C8 spinal cord. Transcranial magnetic stimulation (TMS) was applied to the primary motor cortex (M1) hotspot corresponding to the first dorsal interosseous (FDI) muscle, innervated by the C8 nerve. A 500 ms-duration LIFU was delivered to the C8 spinal cord 400 ms prior to single-pulse TMS over the FDI hotspot. Spinal cord ultrasound stimulation (SCUS) was administered with varying acoustic parameters: intensities (I_SPPA_: 2.5 and 10 W/cm²), DCs (10% and 30%), and PRFs (500 and 1000 Hz). Changes in corticospinal excitability were assessed by comparing TMS-elicited motor-evoked potentials (MEPs) between active and sham SCUS conditions.

**Results:** SCUS with an I_SPPA_ of 10 W/cm², a DC of 30%, and a PRF of 1000 Hz significantly reduced MEP amplitudes compared to sham stimulation. However, at the high intensity (I_SPPA_ of 10 W/cm²), varying the DC between 10% and 30% did not affect MEP amplitudes. Additionally, while a PRF of 1000 Hz decreased MEP amplitudes at 10 W/cm², a PRF of 500 Hz did not produce significant changes.

**Conclusions:** The results indicate that ultrasound stimulation of the spinal cord can suppress corticospinal drive to muscles, especially when utilizing high intensity and high PRF parameters. This suggests that ultrasound stimulation may provide a novel method for modulating human spinal neural activity.

## Introduction

Spinal cord stimulation (SCS) represents a promising method for promoting functional recovery in a range of neurological conditions, such as spinal cord injury, stroke, Parkinson’s disease, and multiple sclerosis [1–9]. SCS functions by modulating spinal neuronal activity and synaptic transmission [10,11], consequently affecting the connections between cerebral and spinal networks and the integration of sensorimotor processes. SCS encompasses two main approaches: epidural SCS (eSCS) and transcutaneous spinal direct current stimulation (tsDCS). eSCS, an invasive method, utilizes implanted electrodes to deliver pulsed currents to the dorsal roots, thereby activating motoneurons through a transsynaptic pathway involving Ia sensory afferents and proprioceptive feedback. This process enables the selective activation of muscles innervated by specific dorsal roots [12,13]. tsDCS, a noninvasive technique, employs surface electrodes to modulate the resting membrane potential of spinal neurons. The effects of tsDCS are determined by the polarity of the applied current: cathodal tsDCS depolarizes neuronal membranes, enhancing spinal Hoffmann reflexes and corticospinal excitability, while anodal tsDCS hyperpolarizes membranes, thereby suppressing these responses [14–17].

Both eSCS and tsDCS present distinct advantages for spinal cord stimulation, yet they also possess inherent limitations. eSCS facilitates selective stimulation of dorsal root ganglia, enabling precise spatiotemporal neuromodulation synchronized with movement [18,19]. However, its invasive nature introduces potential risks such as infection, electrode migration, and spinal compression, which limit its widespread clinical application [20–22]. Conversely, tsDCS, being noninvasive and more accessible, often yields inconsistent outcomes. Some studies have reported no significant polarity-dependent effects, while others have observed predominant anodal facilitation [23,24]. This variability is likely attributable to factors such as neuronal topography, circuit interactions, and electrode configuration [25,26]. Moreover, the limited muscle recruitment selectivity of tsDCS constrains its efficacy in precise motor function restoration [27]. Consequently, there is a pronounced need for the advancement of alternative spinal cord stimulation methodologies.

Low-intensity focused ultrasound (LIFU) is an innovative noninvasive neuromodulation technique offering high spatial precision and deep tissue penetration. LIFU exerts its neuromodulatory effects via mechanical interactions between acoustic waves and mechanosensitive ion channels, inducing both short- and long-term changes in neural circuit activity [28,29]. Its efficacy has been demonstrated in modulating neural activity across various animal species [30–36] and in human brain regions, including the somatosensory, motor, and visual cortices, as well as the thalamus [32,37–44]. With the capability to penetrate soft tissues at wavelengths of approximately 100 μm [45], LIFU can deliver focused energy deep into the spinal cord with high spatial precision. As sound waves travel through soft tissues at an approximate speed of 1.5 km/s, LIFU also achieves remarkable temporal accuracy of less than 1 ms [46]. The high spatiotemporal precision and deep tissue penetration of LIFU make it a compelling alternative for spinal cord stimulation [45–47]. Notably, preclinical studies have shown the feasibility of LIFU for spinal cord neuromodulation, demonstrating its ability to modulate corticospinal pathways, suppress spinal reflexes, and influence motor signal transmission in animal models [48–53]. Despite these promising preclinical findings, the potential of LIFU for modulating neural activity within the human spinal cord remains largely unexplored.

To investigate the effects of LIFU on spinal cord neuromodulation, we combined LIFU with transcranial magnetic stimulation (TMS) to assess its impact on TMS-evoked motor evoked potentials (MEPs), which serve as a measure of corticospinal excitability. A 500 kHz ultrasound transducer, with a focal depth exceeding 100 mm, was employed to precisely target the C8 spinal segment. Single-pulse TMS was delivered to the primary motor cortex (M1) hotspot corresponding to the first dorsal interosseous (FDI) muscle, innervated by the C8 nerve root. A 500 ms ultrasound pulse was applied to the C8 segment 400 ms before the TMS pulse. Spinal cord ultrasound stimulation (SCUS) was administered with varying acoustic parameters: spatial-peak pulse-average intensity (I_SPPA_: 2.5 and 10 W/cm²), duty cycle (DC: 10% and 30%), and pulse repetition frequency (PRF: 500 and 1000 Hz). This study primarily investigated the online effects of SCUS, focusing on changes in TMS-induced MEPs during the SCUS application period. Building on previous findings that suggest an overall inhibitory effect of online LIFU [39,40,54], we hypothesized that SCUS would suppress TMS-evoked MEPs compared to sham conditions in which SCUS was inactive.

## Materials and Methods

### Participants

Thirty-six healthy individuals aged 20 to 30 years participated in the experiment. None of the participants used medications known to affect brain excitability or had any history of neurological or psychiatric disorders. Prior to the experimental sessions, specific details of the experiment and the broader objectives of the study were withheld from participants to maintain research integrity. The study protocol received approval from the Institutional Review Board (IRB) of Texas A&M University, and written informed consent was obtained from each participant before their participation. The study was conducted in compliance with the Declaration of Helsinki.

### Spinal cord ultrasound stimulation (SCUS)

A custom four-element annular array ultrasound transducer (Sonic Concepts Inc., US) operating at a fundamental frequency of 500 kHz was utilized. The transducer features an outer aperture diameter of 64 mm and a curvature radius of 150 mm. The LIFU waveforms were generated using a four-channel Transducer Power Output (TPO) unit provided by Sonic Concepts. The transducer was positioned posteriorly to enable translaminar ultrasound propagation [55], directing the beam through the lamina and soft tissues to target the C8 spinal segment. (Figure 1A). To facilitate accurate positioning, the K-Plan software (Brainbox Ltd., UK), which is a graphical user interface-based ultrasound modeling tool designed for transcranial ultrasound simulation, was employed. Positioning simulations were conducted using data from three human spine X-ray computed tomography (CT) datasets obtained from the open-access VerSe 2020 dataset [56–58]. These simulations enabled the refinement of the transducer placement to ensure optimal ultrasound delivery and precise targeting of the C8 spinal segment. Further details of this simulation-based optimization process are described in the subsequent section. The parameters for SCUS included two spatial-peak pulse-average intensities (I_SPPA_: 2.5 and 10 W/cm²), two duty cycles (DCs: 10% and 30%), and two pulse repetition frequencies (PRFs: 500 and 1000 Hz), as illustrated in Figure 1B.

**Figure 1.**
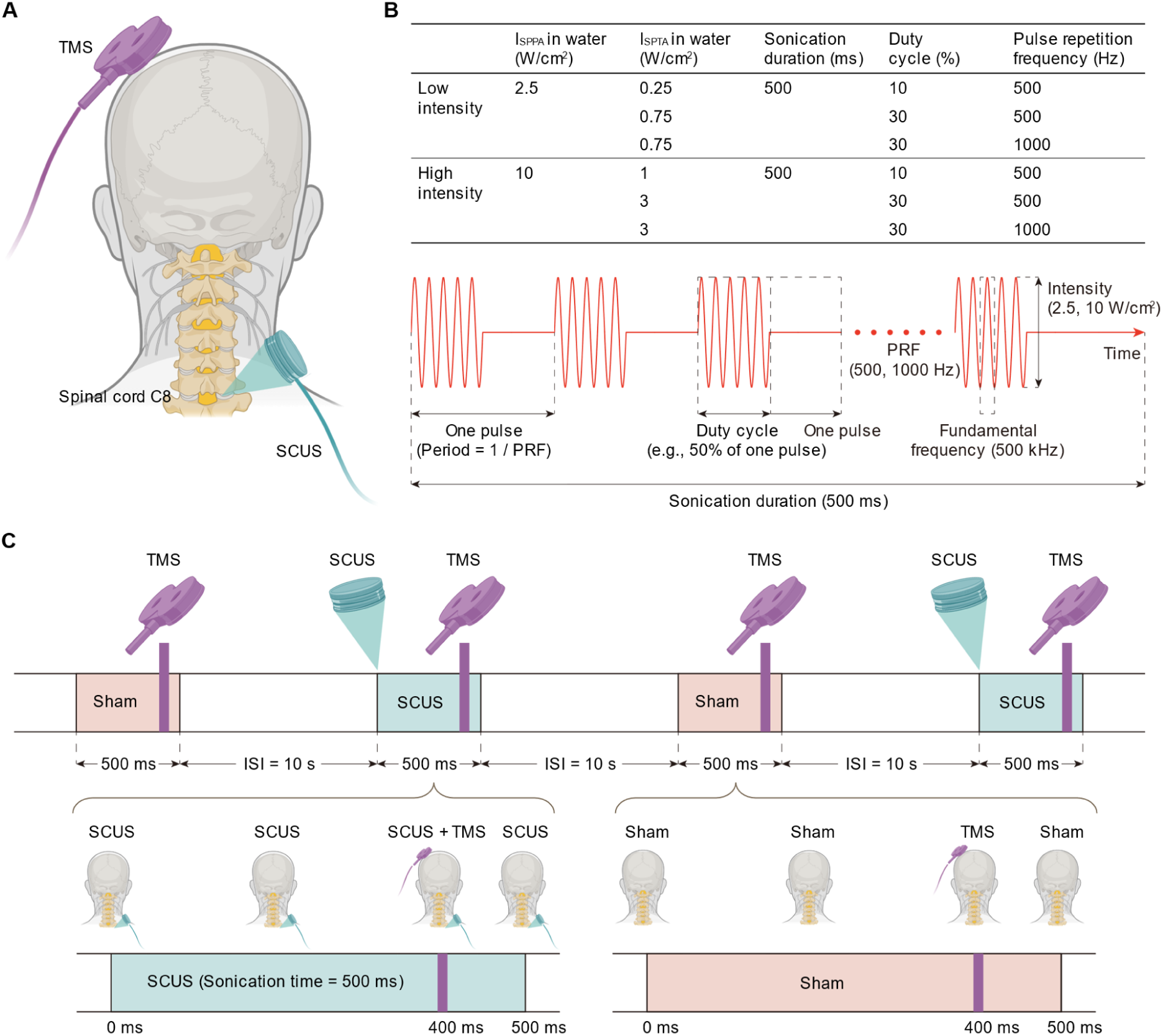
Spinal cord ultrasound stimulation (SCUS) and transcranial magnetic stimulation (TMS). **(A)** Experimental set-up depicting TMS to the primary motor cortex (M1) and SCUS to the C8 spinal cord. **(B)** Acoustic parameters of SCUS that were tested including spatial-peak pulse-average intensity (I_SPPA_: 2.5, 10 W/cm^2^), duty cycle (DC: 10%, 30%), and pulse repetition frequency (PRF: 500, 1000 Hz). The fundamental frequency of 500 kHz and sonication duration of 500 ms were held constant. Eighteen participants with low intensity sonication (I_SPPA_ = 2.5 W/cm^2^) completed three study sessions (DC = 10% and PRF = 500 Hz, DC = 30% and PRF = 500 Hz, DC = 30% and PRF = 1000 Hz). Another group of 18 participants with high intensity sonication (I_SPPA_ = 10 W/cm^2^) completed the same three sessions. **(C)** Schematic illustrating the pulse train level of SCUS in each session. Fifteen sham SCUS conditions and 15 active SCUS conditions were randomized in each session. The inter-stimulus interval (ISI) of 10 s was set between each condition. TMS was delivered 100 ms before the end of sham or active SCUS conditions.

### Transcranial magnetic stimulation (TMS)

To assess changes in corticospinal excitability with or without LIFU applied to the spinal cord, single pulses of transcranial magnetic stimulation (TMS) were administered to the hand representation area of the M1. Motor-evoked potentials (MEPs) elicited by TMS served as indicators of corticospinal excitability during the stimulation period [59]. TMS was delivered using a DuoMAG MP-Dual TMS system (Brainbox Ltd, UK), equipped with a figure-8 butterfly coil featuring two 70 mm windings. The optimal stimulation site, referred to as the hotspot, was identified as the location where the largest MEP was evoked in the FDI muscle with minimal TMS intensity. This hotspot was marked using a frameless neuronavigation system (Brainsight Ltd., Canada) to ensure consistent coil placement during MEP measurements. MEPs were recorded from the FDI muscle using disposable Ag-AgCl surface electrodes (10-mm diameter) affixed to the skin over the muscle belly. The TMS intensity for assessing corticospinal excitability in the FDI was set to evoke MEPs greater than 1 mV in at least 5 out of 10 consecutive trials. MEP signals were amplified and filtered (bandwidth: 30–2000 Hz) using an amplifier (Neurolog System, Digitimer, UK), digitized at a 10 kHz sampling rate via an A/D converter (CED Micro 1401, Cambridge Electronic Design, UK), and stored on a computer for offline analysis.

### SCUS-TMS stimulation delivery

In the SCUS-TMS stimulation protocol, ultrasound transmission gel (Aquasonic 100, USA) was applied between the transducer and the skin to ensure effective energy transfer to the spinal cord. The TMS coil was positioned tangentially to the scalp at a 45° angle to the midline, with the handle oriented laterally and posteriorly to generate a posterior-to-anterior (PA) current in the brain. Single-pulse TMS was delivered at an interstimulus interval (ISI) of 10 seconds. A 500 ms SCUS pulse was applied to the C8 spinal segment, with the TMS pulse synchronized to occur 100 ms before the end of the SCUS pulse. This timing ensured temporal overlap between SCUS and the generation of MEPs induced by TMS, allowing the evaluation of SCUS’s online modulatory effects on corticospinal excitability (Figure 1C). The sham SCUS condition was conducted with the transducer positioned identically to the active condition but without emitting acoustic energy. To ensure blinding, Gaussian white noise was provided to participants during both active and sham conditions to mask auditory cues from the transducer. Each session consisted of 30 SCUS-TMS trials, split into 15 active and 15 sham SCUS trials, randomized within the session to minimize bias. The study included two participant groups. The first group of 18 participants underwent low-intensity sonication (I_SPPA_ = 2.5 W/cm²) across three sessions with different parameter settings: DC = 10% and PRF = 500 Hz; DC = 30% and PRF = 500 Hz; and DC = 30% and PRF = 1000 Hz. The second group of 18 participants received high-intensity sonication (I_SPPA_ = 10 W/cm²) under the same parameter settings. Session order was randomized for each participant, with an ISI of 5 minutes between sessions.

### Ultrasound positioning for spinal cord stimulation

The human cervical spine, composed of stacked, highly irregular vertebrae (Figure 2A), poses significant challenges to ultrasound propagation, leading to aberrations in wavefronts as they travel toward the spinal cord [60]. As ultrasound waves pass through bony structures and surrounding tissues, they undergo absorption, scattering, and refraction, which distort the wavefronts and cause variations in acoustic intensity. To overcome these challenges and optimize ultrasound delivery through the complex anatomy of the cervical spine, the transducer was positioned at a posterior angle, aligned to allow the ultrasound beam to propagate through the lamina of the vertebrae. This approach ensures that the ultrasound beam passes through only a single bone layer, minimizing energy loss and targeting the C8 spinal segment [61]. Specifically, the transducer was placed posterior to the midline on the skin surface and angled obliquely to direct the ultrasound beam through the lamina toward the C8 spinal segment (Figure 2A). This orientation enabled the beam to bypass the central vertebral body and spinous processes, thereby mitigating acoustic interference and attenuation caused by denser bony structures. The beam traveled through the soft tissues and posterior musculature of the neck, maintaining alignment with the translaminar acoustic window, and was focused directly on the C8 spinal cord segment (Figure 2B).

**Figure 2.**
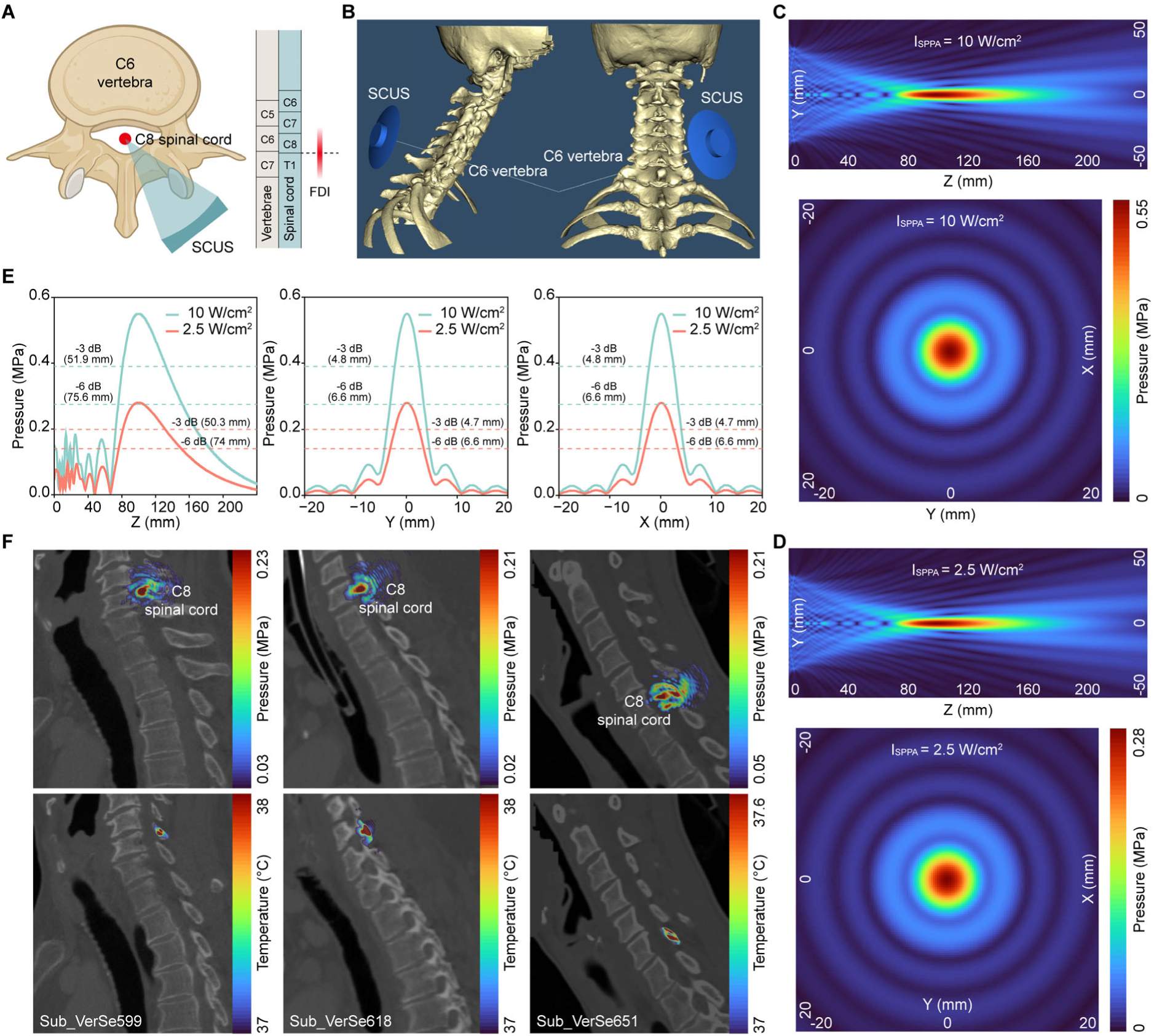
SCUS targeting human C8 spinal cord. **(A)** Ultrasound transducer-C8 spinal cord configuration showing the transducer focused through the right lamina of C6 vertebra. **(B)** Three-dimensional k-Plan viewer depicting position of the ultrasound transducer for C8 spinal cord target. **(C, D)** Ultrasound beam characteristics with focal depth of 100 mm simulated in a free-water field at I_SPPA_ = 10 W/cm^2^ and I_SPPA_ = 2.5 W/cm^2^. Acoustic pressure fields depict the beam in the axial (YZ) plane and the lateral (XY) plane. **(E)** Acoustic pressure profile plots in the free field showing the focal axial length (Z axis) and the focal lateral width (X and Y axes) within the -3 dB and the -6 dB (full width at half maximum, FWHM) areas of the focus. **(F)** SCUS simulations in three human cervical spine computed tomography (CT) data chosen from the VerSe 2020 dataset at the condition of I_SPPA_ = 10 W/cm^2^, DC = 30%, and PRF = 1000 Hz. Acoustic pressure fields overlaid on the CT images demonstrating reliable targeting of the C8 spinal cord. The maximum heating of around 1 °C was found in the posterior arch.

### SCUS simulation

Acoustic models were developed utilizing spinal CT data obtained from the publicly available VerSe 2020 dataset to evaluate SCUS wave propagation through the spinal cord and produce corresponding acoustic pressure maps. Three human spine CT scans of Sub_VerSe599, Sub_VerSe618, and Sub_VerSe651 were chosen due to the complete field of view for C8 spinal cord, sufficient resolution, and no CT artifacts. The transducer was positioned to focus ultrasound through the right lamina of the sixth cervical vertebra to the C8 spinal cord. The positions of the transducer and the spinal cord target were adjusted manually to minimize the intersection of the sonication beam axis with the lamina of vertebral arch. The transducer was located close to the skin surface.

The spinal CT images, defined as the primary planning image in the K-Plan software, were used to generate the spine acoustic and thermal material characteristics. The CT images were resampled according to the spatial size of simulation domain and the number of grid points per wavelength that is defined as 6 in this study. This allows the conversion of CT radiodensity (Hounsfield units, HU) to mass density using the default CT calibration curve defined in K-Plan. The density-based segmentation was implemented to separate CT images into three parts of vertebrae (density threshold of 1150 kg/m^3^), soft tissues (density threshold of 950 kg/m^3^), and background. Within the vertebrae, the bone density was mapped from the CT HU. The sound speed was derived from a density-sound speed linear relationship established in K-Plan. Constant values were set for vertebral absorption coefficient, absorption power law, thermal conductivity, and specific heat. For soft tissues and background, all acoustic and thermal properties were assumed to be constant. Simulations were conducted in the time domain using k-Plan, with acoustic pressure amplitude and phase extracted after the pressure field reached a steady state.

### Data analysis

MEP amplitudes were recorded as peak-to-peak responses using Signal software (Cambridge Electronic Design, UK). The facilitation or inhibition effects of SCUS on MEPs were quantified by normalizing the MEP amplitudes in the active condition as a percentage of those in the sham condition. A percentage below 100% indicates that the conditioning SCUS suppressed the MEP response elicited by TMS, while a percentage above 100% indicates that the conditioning SCUS enhanced the MEP response elicited by TMS. For each sonication intensity condition (I_SPPA_: 2.5 and 10 W/cm²), a repeated-measures ANOVA was conducted to examine the effects of CONDITION (active SCUS vs. sham SCUS) and PARAMETERS (variations in DC and PRF) on normalized MEP amplitudes. The Shapiro-Wilk test was used to assess the normality of the data, while Levene’s test and Mauchly’s test of sphericity were applied to evaluate homogeneity of variances and sphericity, respectively. If the data violated the assumption of normality, a logarithmic transformation was applied. In cases where the assumption of sphericity was violated, the Greenhouse-Geisser correction was used to adjust the analysis.

## Results

### Ultrasound field characterization using free-field simulations

We simulated the three-dimensional acoustic pressure distribution generated by the transducer in a free field for two intensities (I_SPPA_: 2.5 and 10 W/cm²) at a focal depth of 100 mm. This depth provides sufficient range for targeting the C8 spinal cord, even in individuals with larger or thicker neck structures, while enabling efficient beam convergence without premature energy dissipation in superficial tissues. Furthermore, the lamina’s dense bony composition and the acoustic impedance of the posterior musculature are effectively addressed by the extended focal length. This ensures the beam maintains focus as it traverses varying tissue densities, minimizing refraction and scattering caused by irregular bony surfaces. At a target I_SPPA_ of 10 W/cm², the maximum pressure at the focus was 0.55 MPa, with a mechanical index (MI) of 0.78 and a spatial-peak time-average intensity (I_SPTA_) of 1 W/cm² prior to transmission through the spinal cord (Figure 2C). Similarly, at a target I_SPPA_ of 2.5 W/cm², the maximum pressure at the focus was 0.28 MPa, the MI was 0.39, and the I_SPTA_ was 0.25 W/cm² (Figure 2D). Simulations revealed that for an ultrasound beam at I_SPPA_ = 10 W/cm², the -3 dB and -6 dB (full width at half maximum, FWHM) focal axial lengths were 51.9 mm and 75.6 mm, respectively, with corresponding focal lateral widths of 4.8 mm and 6.6 mm (Figure 2E). For an ultrasound beam at I_SPPA_ = 2.5 W/cm², the -3 dB and -6 dB focal axial lengths were 50.3 mm and 74 mm, respectively, with focal lateral widths of 4.7 mm and 6.6 mm (Figure 2E).

### SCUS simulation

Figure 2F (upper panels) displays the simulated ultrasound pressure fields superimposed on CT scans of three human spines—Sub_VerSe599, Sub_VerSe618, and Sub_VerSe651—obtained from the publicly available VerSe 2020 dataset. These simulations demonstrate successful targeting of the C8 spinal cord under two intensity conditions. In the 2.5 W/cm² condition, the pressure at the C8 spinal cord was estimated to be 0.11 ± 0.01 MPa, with a range of 0.11–0.12 MPa. The corresponding I_SPPA_ was 0.39 ± 0.04 W/cm² (range: 0.37–0.44 W/cm²), and the mechanical index (MI) was 0.35 ± 0.07 (range: 0.28–0.42). In the 10 W/cm² condition, the pressure at the C8 spinal cord increased to 0.22 ± 0.01 MPa, with a range of 0.21–0.23 MPa. The I_SPPA_ rose to 1.57 ± 0.17 W/cm² (range: 1.47–1.76 W/cm²), and the MI reached 0.70 ± 0.14 (range: 0.55– 0.83). The estimated attenuation of both focal intensities (I_SPPA_: 2.5 and 10 W/cm²) was approximately 84% on average. The I_SPPA_ and MI for both intensity levels remained below the safety limits recommended by the United States Food & Drug Administration (US FDA). Figure 2F (bottom panels) illustrates the temperature distributions within the same three spines. The temperature rise was predominantly localized in the vertebrae, with average temperatures reaching approximately 38 °C, further confirming the safety of the ultrasound parameters used.

### Estimated temperature rise and safety

Figure 3A depicts temperature changes over time in three individuals under the sonication parameters of I_SPPA_ = 10 W/cm², DC = 30%, and PRF = 1000 Hz, representing the parameter set with the highest potential thermal load. The peak temperature within the vertebrae remained consistently below 38 °C, with estimated temperature rises of 1 °C, 0.97 °C, and 0.55 °C for the three individuals, respectively (Figure 3A). These temperature rises are significantly below the safety threshold of 2 °C established by the Consensus on Biophysical Safety for Transcranial Ultrasonic Stimulation [62], indicating that the thermal risk under these conditions is minimal. Figure 3B illustrates the free-field I_SPTA_ for four different sonication parameter combinations, categorized by intensity and duty cycle (DC): high intensity (HI) with 30% DC, high intensity (HI) with 10% DC, low intensity (LI) with 30% DC, and low intensity (LI) with 10% DC. The corresponding I_SPTA_ values were 3 W/cm², 1 W/cm², 0.75 W/cm², and 0.25 W/cm², respectively. Notably, all free-field I_SPTA_ values in this study remained below the threshold for potential spinal cord damage, as determined by the fitted damage threshold line derived from preclinical studies [63,64]. These results emphasize that the SCUS protocol used in this study adheres to established safety standards.

**Figure 3.**
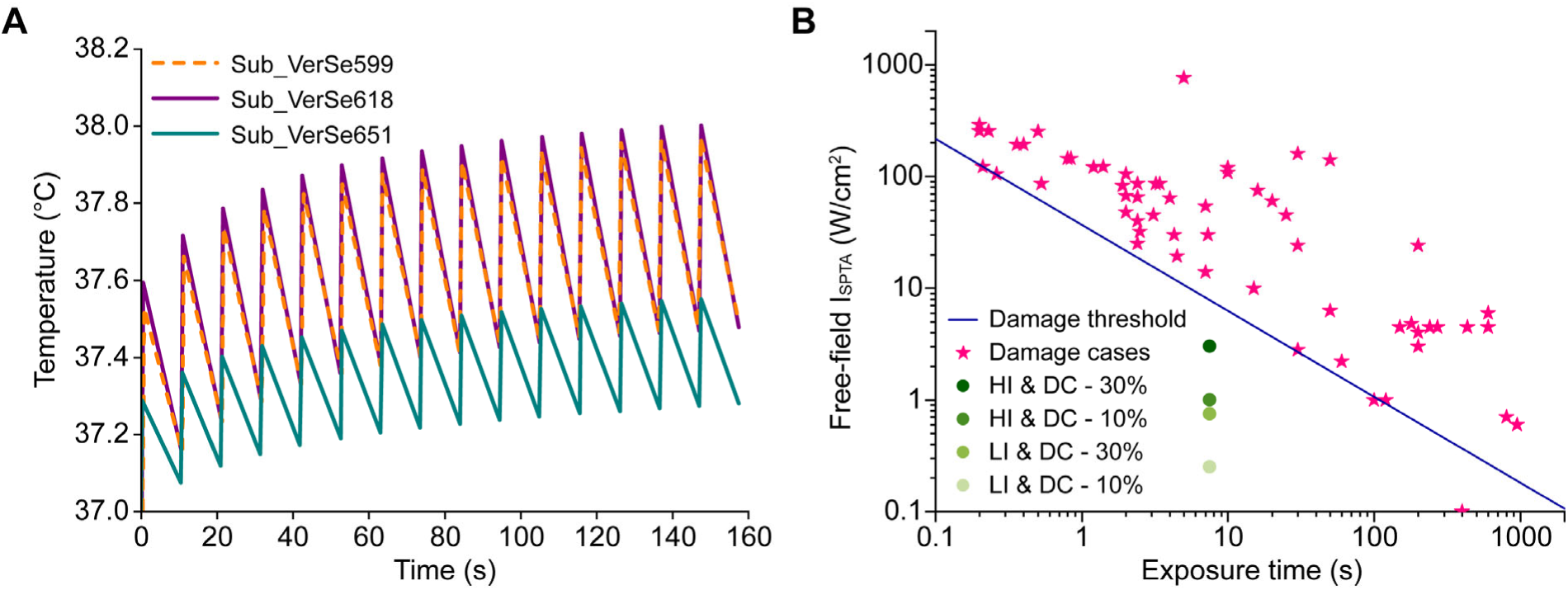
SCUS safety evaluation. **(A)** Temperature changes over time in the three representative individuals at the condition of I_SPPA_ = 10 W/cm^2^, DC = 30%, and PRF = 1000 Hz. The peak temperature rises were 1, 0.97, and 0.55 °C, respectively. The thermal risk is nonsignificant because the temperature rises were lower than 2 °C (Aubry et al., 2023). **(B)** Free-field spatial-peak time-average intensities (I_SPTA_) of four sonication parameter combinations from high intensity (HI), low intensity (LI), 10% DC, and 30% DC were 3, 1, 0.75, and 0.25 W/cm^2^, respectively. All free-field I_SPTA_ values in this study were below the fitted possible spinal cord damage threshold line obtained from pre-clinical cases (Xu et al., 2024a, Xu et al., 2024b).

### Effects of SCUS on corticospinal excitability

Figure 4A depicts changes in MEP amplitude in two representative participants under sham and SCUS conditions. Notably, high-intensity SCUS (I_SPPA_ of 10 W/cm²), with a duty cycle (DC) of 30% and pulse repetition frequency (PRF) of 1000 Hz, resulted in a suppression of mean MEP amplitude compared to sham conditions. In contrast, MEP amplitudes remained relatively stable under low-intensity SCUS conditions (I_SPPA_ of 2.5 W/cm²), with a DC of 10% and PRF of 500 Hz. Figures 4B and 4C display group data illustrating MEP amplitudes during sham and SCUS conditions across different DC and PRF parameters. Figure 4B presents results for low-intensity SCUS (I_SPPA_ of 2.5 W/cm²), while Figure 4C illustrates findings for high-intensity SCUS (I_SPPA_ of 10 W/cm²). For low-intensity SCUS (I_SPPA_ of 2.5 W/cm²), repeated-measures ANOVA revealed no significant effect of CONDITION (F(1,34) = 1.0, *P* = 0.3) and no significant interaction between CONDITION and PARAMETERS (F(2,68) = 0.2, *P* = 0.8) on MEP amplitudes (Figure 4B). Conversely, for high-intensity SCUS (I_SPPA_ of 10 W/cm²), repeated-measures ANOVA demonstrated a significant effect of CONDITION (F(1,34) = 6.7, *P* = 0.01) and a significant interaction between CONDITION and PARAMETERS (F(2,68) = 6.1, *P* = 0.004) on MEP amplitudes (Figure 4C). Post hoc comparisons revealed that for high-intensity SCUS (I_SPPA_ of 10 W/cm²), the MEP amplitude significantly decreased to 78.2 ± 18.6% in active SCUS compared to sham SCUS with the parameter set of DC of 30% and PRF of 1000 Hz (*P* < 0.001). However, no significant differences in MEP amplitude were observed between active and sham SCUS with the parameter sets of DC of 10% and PRF of 500 Hz (*P* = 0.6) or DC of 30% and PRF of 500 Hz (*P* = 0.8). In addition, the suppression in MEP amplitude due to active SCUS were significantly greater with the parameter set of DC of 30% and PRF of 1000 Hz compared to the parameter sets of DC of 10% and PRF of 500 Hz (*P* < 0.001) or DC of 30% and PRF of 500 Hz (*P* = 0.01). These findings suggest that spinal cord ultrasound stimulation can suppress corticospinal excitability (as measured by MEP), especially when employing high-intensity parameters with a high PRF.

**Figure 4.**
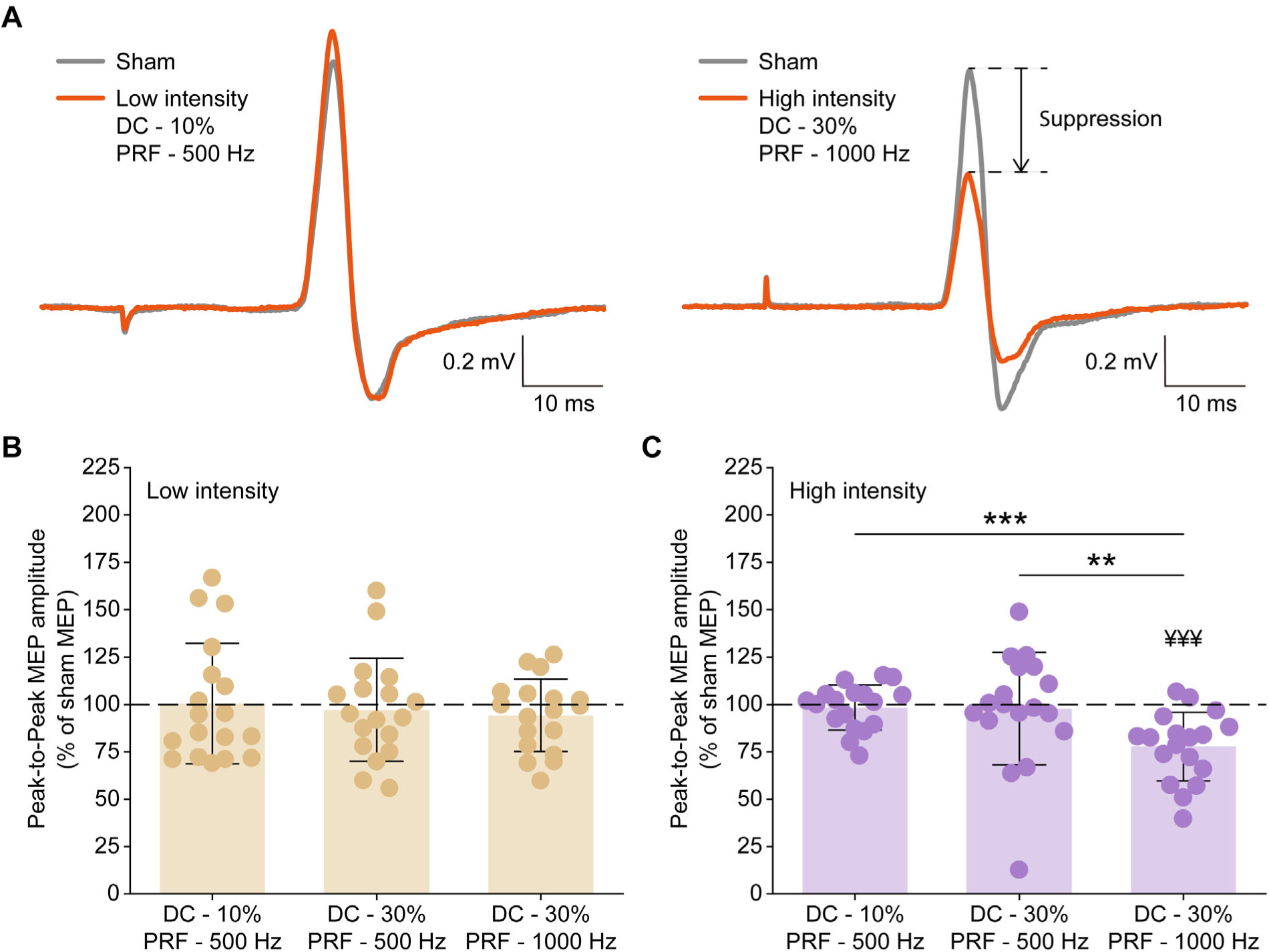
Effect of SCUS on single-pulse TMS-induced MEP amplitudes of the FDI muscle. **(A)** Comparison of mean MEP waveform between sham and SCUS conditions from two representative individuals. High-intensity SCUS (I_SPPA_ of 10 W/cm²), with DC of 30% and PRF of 1000 Hz, suppressed mean MEP amplitude compared to sham condition. **(B)** Group data (*N* = 18) showing low-intensity SCUS had no significant effect on peak-to-peak MEP amplitude. **(C)** Group data (*N* = 18) showing high-intensity SCUS, with DC of 30% and PRF of 1000 Hz, significantly suppressed MEP amplitude compared to sham condition. Peak-to-peak MEP amplitudes of SCUS conditions were normalized to sham MEP amplitudes. The horizontal dotted line represents the MEP amplitude of sham condition. ^¥¥¥^*P* < 0.001, comparison between SCUS MEP and sham MEP. \*\**P* < 0.01 and \*\*\**P* < 0.001, comparison between MEP amplitudes of SCUS with different parameter sets.

## Discussion

Low-intensity focused ultrasound (LIFU) is a promising noninvasive technique for neuromodulation. While focused ultrasound stimulation of the spinal cord has been demonstrated in small animal models [50–52,64], this study represents, to our knowledge, the first investigation of LIFU targeting the human spinal cord to modulate neural activity. The study evaluated the effects of spinal cord ultrasound stimulation (SCUS) on corticospinal excitability, assessed through motor-evoked potentials (MEPs) elicited by transcranial magnetic stimulation (TMS). The findings indicate that SCUS can modulate corticospinal pathways in a parameter-dependent manner, with high-intensity sonication and high pulse repetition frequency (PRF) producing significant suppressive effects on MEP amplitudes. These results highlight focused ultrasound as an effective and noninvasive approach for neuromodulation of the human spinal cord.

Simulations of SCUS targeting the C8 spinal cord were conducted to assess heat and pressure distribution. The results demonstrated that temperature increases caused by SCUS were predominantly localized to the vertebrae, with rises within the C8 spinal cord tissue remaining below 1 °C. This minimal and localized heating effect is attributed to the focused ultrasound beam and the energy dissipation properties of the vertebrae and surrounding soft tissues. As dense structures, the vertebrae absorb and dissipate ultrasound energy, preventing significant thermal buildup in the spinal cord. Notably, these temperature increases were well below the established safety threshold of a 2 °C rise, as outlined by the Consensus on Biophysical Safety for Transcranial Ultrasonic Stimulation [62]. Furthermore, the simulated mechanical index (MI) values were significantly below the US FDA safety limit (MI < 1.9), with high-intensity SCUS yielding an MI of 0.70 ± 0.14 and low-intensity SCUS producing an MI of 0.35 ± 0.07. The approximately 84% attenuation of focal intensity through the bony and soft tissue structures of the cervical spine, coupled with the fact that all free-field I_SPTA_ values remained below preclinical damage thresholds (Figure 3B) [63,64], further supports the minimal thermal and mechanical risks associated with SCUS.

Our study investigated the online effects of SCUS, specifically examining changes in TMS-induced MEPs during SCUS application. The findings reveal that high-intensity SCUS (I_SPPA_ of 10 W/cm²) significantly suppresses MEP amplitudes when applied with sonication parameters of a 30% duty cycle (DC) and a pulse repetition frequency (PRF) of 1000 Hz. This reduction in MEP amplitudes indicates that high-intensity SCUS inhibits corticospinal excitability. In contrast, low-intensity SCUS (I_SPPA_ of 2.5 W/cm²) did not result in any significant changes in MEP amplitudes, regardless of the DC or PRF settings. Our experimental approach parallels previous studies investigating the online effects of LIFU over the primary motor cortex (M1) on MEP amplitudes [40,54,65]. In these studies, an ultrasound transducer attached to a TMS coil was used for concurrent stimulation to assess online effects of LIFU. Consistent with our findings, these studies demonstrated that LIFU applied over M1 attenuates MEP amplitudes [40,54,65,66]. Previous research has highlighted the parameter-dependent nature of LIFU-induced changes in TMS-evoked MEP amplitudes [54,67]. In agreement with our results showing that SCUS with a 30% DC and a PRF of 1000 Hz inhibits corticospinal excitability, earlier studies have reported similar effects. Specifically, LIFU applied to M1 using comparable parameters— PRF of 1000 Hz and DC of 36% or 30%—was shown to suppress corticospinal excitability [40,68]. Additionally, our results highlight the importance of sonication intensity in achieving effective neuromodulation. Only high-intensity SCUS (I_SPPA_ of 10 W/cm²) produced significant modulation of MEP amplitudes, suggesting that the total energy delivered is a key determinant of SCUS-induced neuromodulatory effects. Interestingly, Ennasr et al. reported a stronger inhibitory effect with lower-intensity LIFU applied to M1 compared to higher-intensity LIFU (I_SPPA_ of 6 W/cm² vs. 24 W/cm²) [65]. However, their study involved significantly less transcranial attenuation (∼69%) than the attenuation encountered in our spinal cord targeting (∼84%). This substantial difference in attenuation implies that the energy delivered to the C8 spinal segment under our low-intensity SCUS condition (I_SPPA_ of 2.5 W/cm²) may have been insufficient to elicit a significant modulatory effect.

The mechanisms through which ultrasound stimulation modulates neural activity remain not fully understood but are hypothesized to involve various physical interactions with biological tissues, such as mechanical forces, localized thermal effects, and auditory stimuli [46]. The mechanical pressure generated by SCUS can directly activate mechanosensitive TRAAK channels [69,70]. These channels inhibit neuronal activity by mediating potassium efflux, leading to membrane hyperpolarization and decreased spinal excitability [69]. This reduction in spinal excitability is likely to lead to a decreased recruitment of alpha motor neurons, resulting in diminished MEP amplitudes elicited by TMS. Our findings are unlikely to be attributed to thermal effects, as simulations of SCUS-induced temperature changes in the C8 spinal cord showed a minimal increase (< 1 °C), significantly below the established threshold for thermal bioeffects [32,38,42]. To mitigate potential auditory confounds associated with ultrasound stimulation, particularly at a PRF of 1000 Hz [71], Gaussian white noise masking was utilized throughout the SCUS experiments. Participants were consistently exposed to auditory stimuli, including transducer noise and masking noise, across all SCUS sessions. Significant modulation of MEPs occurred exclusively under specific sonication parameters, indicating a parameter-dependent response. This strongly suggests that the observed MEP changes were directly attributable to SCUS rather than general auditory stimulation. Consequently, we propose that auditory confounds had a minimal impact on our findings.

SCUS’s capability to suppress corticospinal excitability holds therapeutic potential. Disorders characterized by excessive excitability of the corticospinal or spinal circuits, such as spasticity and hyperekplexia, could benefit from targeted SCUS interventions. By optimizing sonication parameters, clinicians could selectively modulate specific spinal segments, offering a noninvasive alternative to invasive neuromodulation techniques like epidural spinal cord stimulation. The parameter-dependent nature of SCUS effects also highlights its versatility as a neuromodulation tool. While our findings demonstrated inhibitory effects on corticospinal excitability, alternative sonication parameters could elicit excitatory responses [72–74], broadening the range of applications for SCUS. The potential for SCUS to induce offline effects—sustained modulation of corticospinal excitability following stimulation—offers further opportunities for therapeutic development, including interventions targeting neural plasticity. Notably, the favorable safety profile demonstrated in this study, with negligible thermal and mechanical risks, reinforces the feasibility of SCUS for clinical translation.

This study has several limitations that warrant consideration. First, the simulations included data from only three subjects, which may not fully capture the variability in spinal morphology or the range of acoustic and thermal responses to sonication across diverse populations. The accuracy of our SCUS simulations may be limited by uncertainties in the acoustic properties of the vertebrae and surrounding soft tissues, including variations in attenuation, absorption, and speed of sound. These parameters were estimated based on literature values and may not fully capture the complex interactions of ultrasound with human spinal cord. Second, our study only assessed a limited range of sonication parameters due to practical constraints, such as study time, and safety concerns regarding potential thermal bioeffects from prolonged sonication. Although the sonication duration was fixed at 500 ms in this study, future research should explore how varying duration affects SCUS outcomes and its interactions with other parameters, including intensity, duty cycle, and pulse repetition frequency. Previous studies have demonstrated that sonication duration significantly affects the outcomes of LIFU [54,74], highlighting the need for further investigation. Third, our study design raises the potential concern of carry-over effects between SCUS sessions. While previous studies utilizing concurrent ultrasound stimulation and TMS have not found evidence of cumulative effects [39,75], additional measures were implemented to address this possibility. The order of SCUS sessions was counterbalanced across participants, and rest intervals were included between sessions to reduce the likelihood of residual effects from earlier sonication.

In conclusion, this study demonstrates that the spinal cord ultrasound stimulation (SCUS) modulates corticospinal excitability in a parameter-dependent manner, with high-intensity sonication under specific duty cycle and pulse repetition frequency conditions significantly suppressing MEP amplitudes. These results highlight the potential of SCUS as a noninvasive neuromodulation technique to modulate spinal circuits. Future research should focus on optimizing sonication parameters, exploring long-term effects, and evaluating SCUS in clinical populations to fully realize its therapeutic potential.

## Data availability statement

The data that support the findings of this study are available from the corresponding author upon reasonable request.

